# Epigenetics and expression of key genes associated with cardiac fibrosis: *NLRP3, MMP2, MMP9, CCN2/CTGF*, and *AGT*

**DOI:** 10.1101/2020.10.23.352518

**Authors:** Sruti Chandra, Kenneth C. Ehrlich, Michelle Lacey, Carl Baribault, Melanie Ehrlich

## Abstract

Excessive inflammatory signaling and pathological remodeling of the extracellular matrix are important contributors to cardiac fibrosis and involve major changes in gene expression. We examined the relationships between tissue-specific expression and the epigenetics of five genes involved in these pathways, *NLRP3, MMP2, MMP9, CCN2/CTGF,* and *AGT.* The proteins encoded by these genes play major fibrosis-related roles in inflammasome formation, degradation of extracellular matrix proteins and remodeling of the extracellular matrix and vasculature, autocrine regulation of fibrosis, or cell signaling. Our analyses showed that the first four of these genes had super-enhancers (unusually strong enhancer clusters) that correlate with their very high expression in monocytes, neutrophils, fibroblasts, or venous cells. Expression of the gene encoding miR-223, a micro-RNA that plays an important role in downregulating NLRP3 protein levels, is also probably driven by the super-enhancer in which it is embedded. Enhancer chromatin for all these genes was inside as well as outside the gene body. While *AGT,* which encodes precursors of angiotensin II, lacked a super-enhancer, its tissue-specific expression profile correlates with the tissue-specific enhancer chromatin extending into its distant silent gene neighbor (*CAPN9*). Tissue-specific peaks of DNA hypomethylation, open chromatin (DNaseI hypersensitivity), and transcription factor binding were detected in subregions of these super-enhancers/enhancers that are likely to be the main drivers of expression of their associated gene. We found that *CCN2/CTGF* is co-expressed with its far-upstream neighbor *LINC01013*, a noncoding RNA gene, specifically in vein endothelial cells (HUVEC) and chondrocytes. Evidence from chromatin looping profiling (Hi-C) suggests coregulation of these genes in HUVEC. Our findings indicate the importance of understanding the often-overlooked roles of enhancers and their hypomethylated, transcription factor-binding subregions in the regulation of expression of fibrosis-related genes in normal and fibrotic tissue.

## 1. Introduction

Fibrotic disease is estimated to be responsible for almost half of the deaths in the US [1], and its far-reaching consequences extend to COVID-19’s effects on disparate tissues and organs [2]. Cardiac fibrosis involves the remodeling of the heart accompanied by the excessive accumulation of extracellular matrix (ECM) proteins due to chronic stress, injury, systemic disease, or drugs [3, 4]. Because the scar tissue impedes the ability of the heart to pump properly, cardiac fibrosis plays a causative role in many types of heart disease, including atrial fibrillation, heart failure, and impaired systolic and diastolic function. The main cell types that contribute to cardiac fibrosis include cardiac fibroblasts (which constitute ~70% of the nuclei in the heart), myofibroblasts (which very highly express ECM proteins and are derived from cardiac fibroblasts or infiltrating fibrocytes), cardiomyocytes, cardiac endothelial cells, monocytes, neutrophils, and T cells [4, 5].

Epigenetics refers to the inherited changes from cell to cell that involve DNA and chromatin but do not alter the sequence of bases in DNA. Changes in epigenetics, usually histone or DNA modifications, and alterations in the composition and conformation of chromatin provide a kind of cellular memory. This memory is needed for differentiation and, to a lesser extent, for sustained responses to some types of physiological changes [6], including in the heart [7]. Epigenetics is involved in turning genes on or off, up- or down-modulating their levels of expression, or maintaining levels of transcription or transcription repression in normal and diseased tissues [6, 8]. The importance of understanding the normal epigenetic modifications of cardiac fibrosis-associated genes is underscored by the involvement of differentiation in fibrosis, namely, transdifferentiation of fibroblasts and fibrocytes to myofibroblasts and inflammation-associated differentiation of monocytes to macrophages [9].

The critical role of epigenetic changes in cardiac fibrosis is beginning to be elucidated [7, 9–12]. For example, Williams et al. [3] examined the involvement of histone acetylation/deacetylation, the most extensively studied type of histone modification involved in transcription control, in a mouse model of cardiac fibrosis. They found that the inhibition of histone deacetylase (HDAC) activity targeted to one subclass of HDACs (class I) suppressed cardiac fibrosis. However, the effects of HDACs on cardiac fibrosis can be indirect. They can reflect histone acetylation changes that affect cell signaling or transcription factor (TF) activity rather than directly targeting the gene of interest, and they can involve acetylation of non-histone proteins [4, 13]. In addition to chromatin epigenetic analyses, disease- and transcription-associated changes in DNA methylation were found in the promoter regions of fibrosis- and inflammation-related genes [4, 14, 15].

Among the major contributors to cardiac fibrosis and related heart disease are the deleterious activation of the autocrine/paracrine renin-angiotensin system (RAS), excessive inflammatory signaling from infiltrating monocyte/macrophages, and pathological ECM remodeling upon upregulation and transdifferentiation of cardiac fibroblasts or infiltrating fibrocytes [16–18]. We chose genes from each of these pathways to study relationships between tissue-specific epigenetics and transcription and to identify regions in the vicinity of these genes that are candidate cis-acting elements contributing to gene dysregulation in cardiac fibrosis. These genes are *AGT* (Angiotensinogen), *NLRP3* (NLR Family Pyrin Domain Containing 3)*, CCN2/CTGF* (Cellular Communication Network Factor 2/Connective Tissue Growth Factor), and *MMP2* and *MMP9* (Matrix Metallopeptidase 3 and 9). *AGT* encodes the precursors of angiotensin II (Ang II), a fibrosis-inducing growth factor and signaling molecule as well as an endocrine vasoconstrictor [19], which can be expressed by myofibroblasts as well as by liver and other organs [17]. Ang II is on one of the pathways for priming and activating NLRP3 inflammasome formation [18] as well as being a key stimulus for cardiac fibrosis [19]. Activation of NLRP3 inflammasomes, which involves upregulation of *NLRP3* gene expression, results in the production of IL-1β and IL-18, which, in turn, promotes myofibroblast formation, and can also initiate pyroptosis to induce cardiac inflammation. NLRP3 is also critical for inflammation associated with many other diseases (including cancer, colitis, gout, type II diabetes, and neonatal onset multisystem inflammatory disease) as well as its normal protective roles in responding to a wide range of damage signals [20]. NLRP3 protein can also contribute to fibrosis by favoring the epithelial-to-mesenchymal transition in both inflammasome-dependent and independent pathways [21]. In addition, abnormal NLRP3 inflammasome signaling can lead to arrhythmic sarcoplasmic reticulum Ca^+2^ release in the heart promoting atrial fibrillation [18]. CTGF, the product of the *CCN2/CTGF* gene, is an autocrine regulator of fibrosis [16] with many disparate cell type-specific effects including the induction of angiogenesis [22, 23]. It upregulates cardiac fibrosis through its stimulation of cardiac fibroblasts although it is also upregulated in injured cardiomyocytes [16]. MMP2 and MMP9 are closely related type IV collagenases that help shape the ECM and may provoke or ameliorate cardiac fibrosis depending on the context [5].

Despite the importance of these genes to cardiac fibrosis and the major role of enhancers (chromatin that upregulates, in *cis*, promoters at the 5’ end of genes and is often distant from the promoter) in tissue-specific gene expression [6], analysis of the epigenetics of these genes has been mostly limited to promoter regions [14, 23–27]. However, there have been notable exceptions [14, 25, 28, 29]. In the present study we describe important epigenetic features within gene bodies and in the gene neighborhoods of *NLRP3, MMP2, MMP9, CCN2/CTGF,* and *AGT* as well as non-coding RNAs whose dysregulation might contribute to abnormal expression of these genes in disease.

## 2. Materials and methods

### 2.1 Transcriptomics

RNA-seq data for tissues and skin fibroblast cell lines were obtained from the Genotype-Tissue Expression database (GTEx) using median gene expression levels derived from hundreds of samples for each tissue type. [30]. For cell cultures, RNA-seq data were from poly(A)^+^ RNA in color-coded overlaid graphs in the figures (ENCODE/Wold Lab at Caltech; vertical viewing range, 0 – 8 except for *AGT* for which it was 0 – 12; UCSC Genome Browser [31]) with quantification of RNA-seq data as previously described [32]. In addition, strand-specific RNA-seq profiles (ENCODE/Cold Spring Harbor Labs[31]) were used to confirm which strand was transcribed and to examine additional cell types (UCSC Genome Browser [31]).

### 2.2 Epigenetics

Epigenetic data were from the Roadmap Epigenetics Project [6] downloaded from UCSC Genome Browser hubs (http://genome.ucsc.edu/) [31]. The 25-state chromatin state segmentation tracks were used except for *CCN2* for which the 18-state tracks were used because they gave fewer uninformative “low-signal” segments. The original color-coding of Roadmap data for chromatin states as shown in the figures was slightly modified to increase clarity. The displayed histone acetylation profiles (vertical viewing range 0 - 10) were the same ones that had been used in determination of chromatin state segmentation [6]. The tissues for chromatin states were from males except for lung, monocytes, and B cells [6, 31] and prefrontal cortex (a mixture of 81-yo male and 75-yo female samples), and cell cultures were previously described [32]. DNaseI hypersensitivity profiling (vertical viewing range 0 – 40) and DNA methylation profiling by bisulfite-seq usually came from the same samples as for the chromatin state [6]. Additional bisulfite-seq profiles for a control and an atherosclerotic aorta sample (Aorta ctl #2, thoracic aorta, and Atheroscl aorta, aortic arch) came from an 88 yo female [33] and are available at the UCSC Genome Browser (93N and 93A, Zaina-2014 at the DNA Methylation Hub). Atherosclerosis-specific and aorta-specific DNA hyper- or hypomethylation was determined from bisulfite-seq data as previously [34]. At the UCSC Genome Browser, the monocytes for the bisulfite-seq track are incorrectly labeled as macrophages. Super-enhancer designations were from the dbSUPER website [35] and checked by inspection of histone acetylation tracks at the UCSC Genome Browser.

### 2.3 Transcription factor binding profiles

A limited number of whole-genome TF binding profiles obtained by chromatin immunoprecipitation/next-gen sequencing (ChIP-seq) on cell types expressing the genes highlighted in this study are publicly available. The ENCODE 3 Transcription Factor ChIP-seq Peaks provided the examined TF binding data (UCSC Genome Browser [31].

## 3. Results

### 3.1 The inflammasome-associated NLRP3 gene has a super-enhancer mirroring its high and specific expression in monocytes and neutrophils

*NLRP3* encodes a protein that is a limiting factor for activation of NLRP3 inflammasome formation [36], which is important in inflammation-related cardiovascular disease [37]. It is expressed at a much higher level in leukocytes (median of 755 samples, 23 TPM (transcripts per kilobase million; GTEx RNA-seq database [30]) than in any of the 49 other examined normal tissues (Fig. 1A). *NLRP3* displayed enhancer chromatin (orange segments) within the gene body and surrounding the gene specifically in monocytes, neutrophils, and lymphocytes (Fig. 2B); macrophage epigenetic data are not available. Enhancer chromatin had been determined from enrichment in histone H3 lysine 27 acetylation (H3K27ac) plus H3 lysine 4 monomethylation (H3K4me1) [6]. Active promoter chromatin (Fig. 2B, red segments) at the transcription start site (TSS; Fig. 2A, broken arrow) was seen in these leukocytes but also in liver, which had very little *NLRP3* RNA and lacked appreciable amounts of enhancer chromatin. Active promoter chromatin had been determined from enrichment in H3K27ac and H3K4me3, instead of the H3K3me1 enrichment that helps define enhancers [6]. Importantly, *NLRP3* did not just display traditional enhancers in monocytes and neutrophils, but rather a super-enhancer, which is an unusually large cluster (>3 kb) of enhancer chromatin regions that can include some promoter chromatin (Fig. 2B, dotted line) [35, 38]. Super-enhancers are typically associated with especially strong, differentiation-related expression of the related gene.

**Fig. 1.**
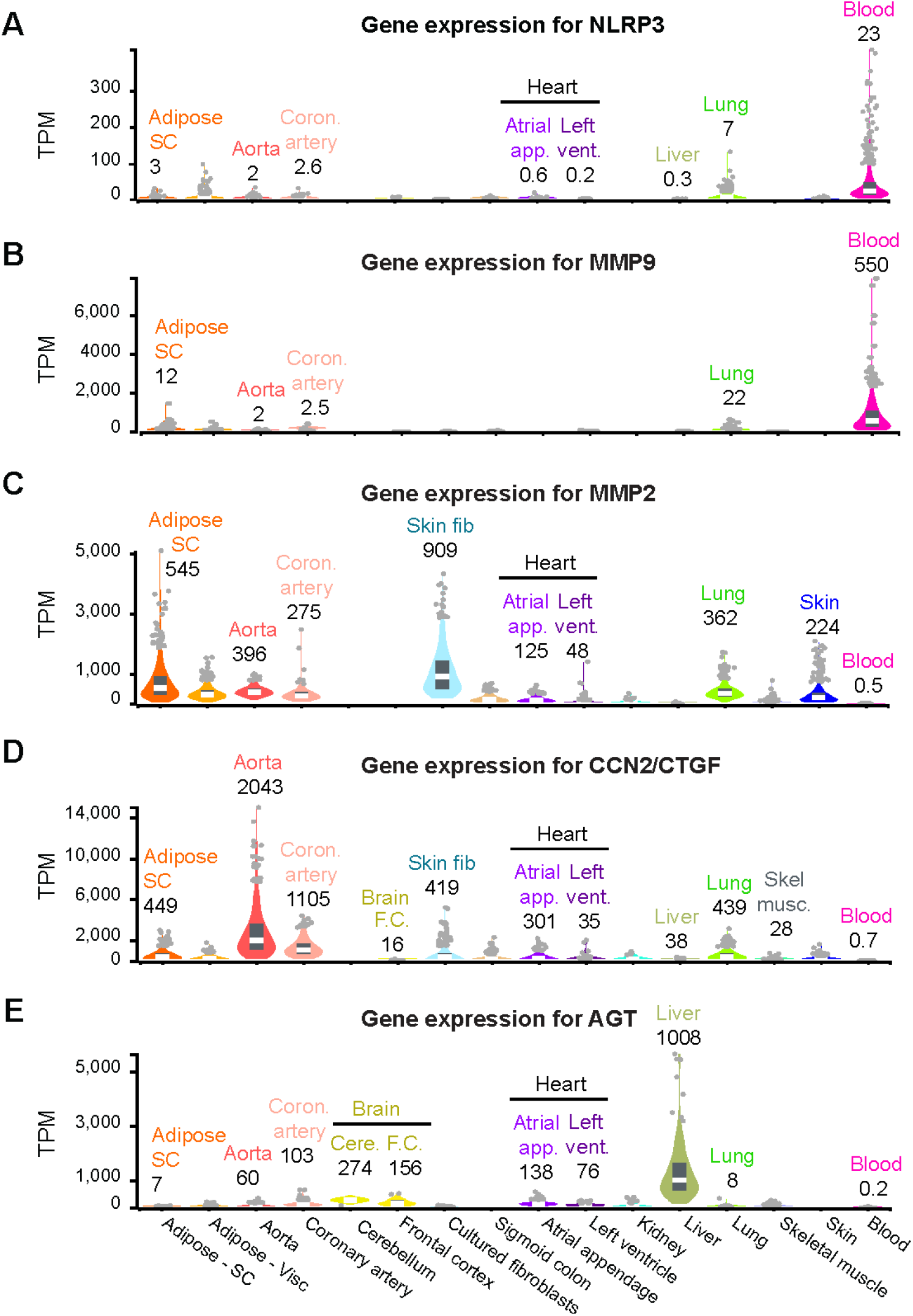
Tissue-specificity of gene expression is similar for *NLRP3* and *MMP9* but dissimilar for *MMP2, CCN2/CTGF,* and *AGT*. RNA-seq data (violin plot) for expression of the indicated genes in 15 tissues and in cultured skin fibroblasts is shown; there were hundreds of biological replicates for each tissue type. Select median values are given. The tissues shown are a subset of the 52 tissues in the GTEx database [30]. Adipose SC, subcutaneous adipose; Coron., coronary; brain FC, brain frontal cortex; cere., cerebellum; atrial app., atrial appendage; vent, ventricle; blood, leukocytes.

**Fig. 2.**
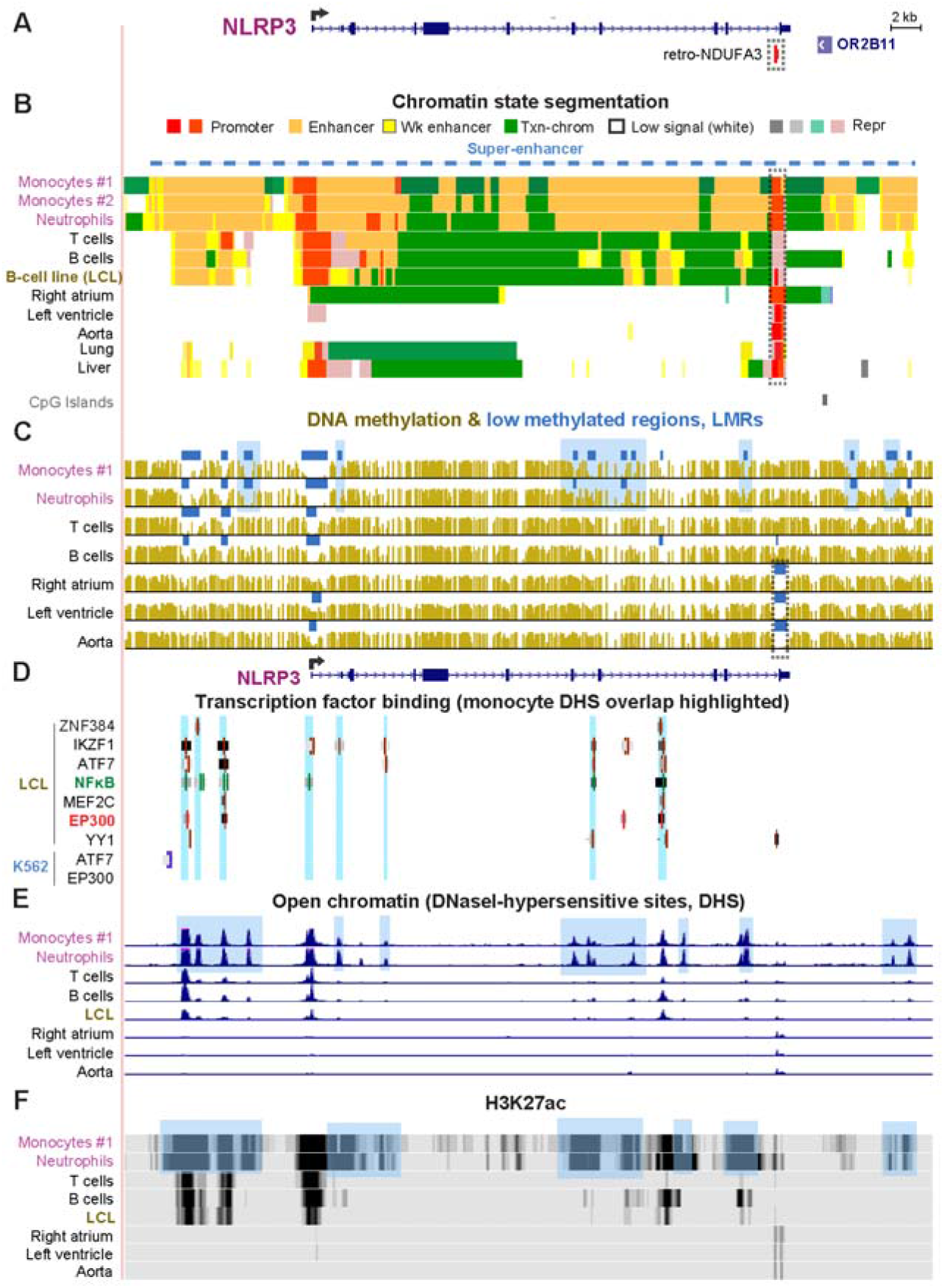
*NLRP3* exhibits a super-enhancer containing foci of DNA hypomethylation and DNaseI hypersensitivity specifically in monocytes and neutrophils, which highly express this gene. (A) The main *NLRP3* (chr1:247,566,540-247,622,212) RefSeq isoform is shown (see Supplemental Fig. S1) and the adjacent olfactory gene *OR2B11,* which is silent in all studied tissues. Broken arrow, the TSS and direction of transcription. (B) Chromatin state segmentation denotes predicted promoter (red), strong enhancer (orange), weak enhancer (yellow), low-signal or repressed (Rep), or H3K36me3-enriched transcribed (green) types of chromatin. Two monocyte samples are shown for the chromatin state analysis. Blue dashed line, super-enhancer in monocytes and neutrophils. (C) DNA methylation profiles (bisulfite-seq); blue bars, regions that were significantly hypomethylated relative to the same genome (LMRs). (D) Local binding of the TFs in the lymphoblastoid cell line (LCL) or in K562 cells. (E) Open chromatin peaks determined as DNaseI-hypersensitive sites (DHS). (F) H3K27ac signal intensity, indicative of the strength of the promoter and enhancer chromatin. Blue highlighting in (C), (E), and (F), monocyte/macrophage-specific epigenetic marks. All tracks are from hg19 in the UCSC Genome Browser and are aligned in this and subsequent figures.

Within the *NLRP3* super-enhancer, there were regions of especially low DNA methylation (low methylated regions, LMRs; blue horizontal bars, Fig. 2C) that were specific to monocytes and/or neutrophils. The monocyte/neutrophil-associated LMRs usually overlapped myeloid-specific regions of open chromatin (DNaseI-hypersensitive sites, DHS) and of high enrichment in H3K27ac (Fig. 2C - E, blue highlighting). A subregion containing 3 CpGs located 0.6 kb upstream of the TSS that was previously shown to be hypomethylated in a monocytic leukemia cell line upon infection with a myobacterium [14] was less methylated in monocytes than in other samples (Supplemental Fig. S1G, blue highlighting).

B cells and T cells, like monocytes and neutrophils, were enriched in promoter chromatin (Fig. 2B) at the *NLRP3* TSS but had much less enhancer chromatin in and around *NLRP3*. B-cells and T-cells were reported to have negligible *NLRP3* expression unless expression was induced, e.g., by lipopolysaccharide [39, 40]. RNA-seq profiles indicated a low level of expression of *NLRP3* in resting B cells and in a B-cell lymphoblastoid cell line (LCL; GM12878) (Supplemental Fig. S1B). Genome-wide profiles of binding by many TFs were available for this LCL and for K562 cells (myelogenous leukemia cell line) [31, 41]. TFs bound in or near open chromatin regions (DHS) in *NLRP3* in the LCL, but not in the analogous regions in *NLRP3*-nonexpressing K562 cells (Fig. 2D and E). These TFs include ZNF384, a regulator of extracellular matrix proteins; IKZF1 (IKAROS), a regulator of hematopoietic cell differentiation; ATF7 and MEF2C, transcriptional activators, EP300, a histone transacetlylase, chromatin remodeling protein, and enhancer/super-enhancer protein; and the RELB subunit of NFκB (Fig. 2D, green lines). NFκB is a signaling TF that is necessary, but not sufficient, for transcriptional activation of *NLRP3* [36]. Many more DHS probably bind EP300 in monocytes and neutrophils than in the weakly *NLRP3*-expressing lymphoblastoid cells; these TF-binding profiles were not available for monocytes or neutrophils.

Surprisingly, we found that the distal (3’) end of *NLRP3* displayed promoter chromatin with the histone marks of active (Fig. 2B, red segments) or poised/repressed (beige segments) promoters in most tissues. This 3’ promoter chromatin (Fig. 2B, dotted rectangle) overlapped a retrotransposon-derived gene, *retro-NDUFA3* near the end of the last intron (Fig. 2A). Only leukocytes had high levels of DNA methylation in this region (Fig. 2C and 10 additional tissue types not shown). These findings suggest that leukocyte-specific DNA methylation repressed most, but not all (Supplemental Fig. S1D), of the 3’ promoter activity that would interfere with *NLRP3* mRNA formation in monocytes and neutrophils. Given the enhancer chromatin adjacent to the 3’ promoter chromatin in monocytes and neutrophils (Fig. 2B), it may be especially important to have local DNA methylation in these cell types. ChIP-seq indicated that the TF YY1, a transcriptional repressor or activator protein, binds within this promoter region (Fig. 2D). Similarly, we had previously demonstrated for *CDH15* and *PITX3* that DNA hypermethylation specific to tissues expressing these two genes repressed an intragenic 3’ promoter [42]. Downstream of *NLRP3* is a gene encoding an olfactory receptor protein (OR2B11) that contains regions of enhancer chromatin in monocytes and neutrophiles despite its negligible expression in all 52 tissues from the GTEx database. Therefore, the enhancer chromatin in the vicinity of *OR2B11* probably upregulates expression of *NLRP3.*

We also examined genes encoding the other three components of NLRP3 inflammasomes. *CASP1* and *PYCARD* (*ASC*) displayed DNA hypomethylation and large amounts of enhancer chromatin specifically in monocytes and neutrophils (*CASP1*) or in leukocytes in general (*PYCARD*; Fig. S2A and B). In contrast, *NEK7* showed less tissue specificity in its expression and epigenetics and had a lower level of RNA in leukocytes than in most other tissues (Fig. S2C). Vento-Tormo et al. [15] examined monocyte-macrophage differentiation-related changes immediately upstream of the promoter regions of *PYCARD* and *CASP1* and found that the differentiation of monocytes was accompanied by DNA hypomethylation in these regions concomitant with gene upregulation. In the methylated subregions that we saw immediately upstream of the promoter, we noted enhancer chromatin in monocytes (Fig. S2A and B). The inferred enhancer might increase in strength upon demethylation during differentiation to macrophages.

### 3.2 The gene encoding miR-223, which downregulates NLRP3 protein production, has a monocyte- and neutrophil-specific super-enhancer, like NLRP3

MicroRNA (miRNA) miR-223, is a critical regulator of NLRP3 protein levels by decreasing *NLRP3* mRNA stability and translation [43]. Its host gene, *hsa-mir-223*, is transcribed preferentially in leukocytes as well as in spleen (TPM, 185 and 146, respectively) unlike *NLRP3* which is transcribed preferentially only in leukocytes (TPM, 23 and 6, respectively). We found that the 5-kb *hsa-mir-223* gene is embedded in a 38-kb super-enhancer seen specifically in monocytes and neutrophils (Fig. 3D, dashed line). The super-enhancer overlapped DHS and pockets of DNA hypomethylation that were most prominent in monocytes and neutrophils (Fig. 3E and G, blue highlighting) and stopped at the end of the neighboring *VSIG4* gene, which has a different tissue specificity.

**Fig. 3.**
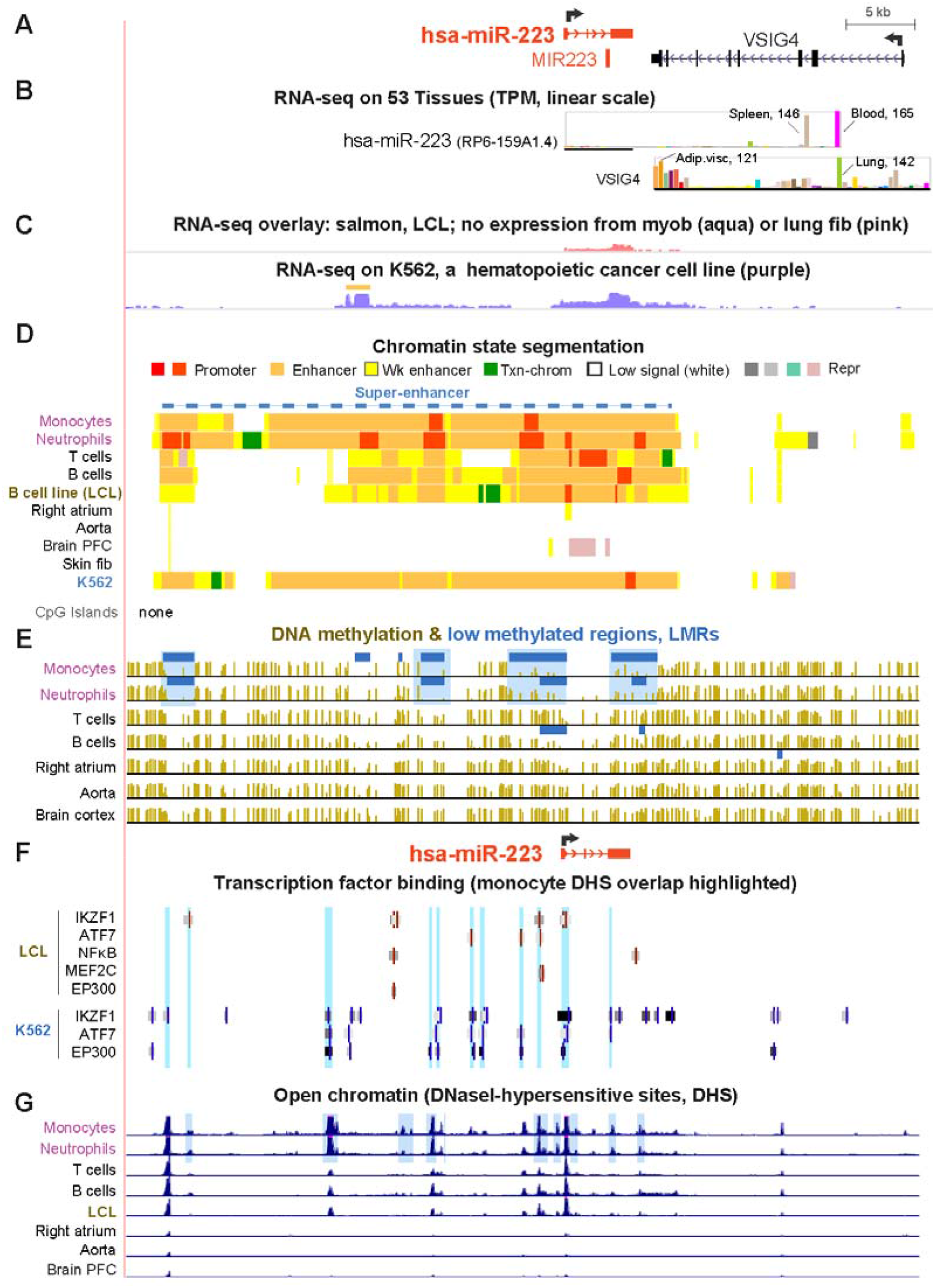
The host gene encoding the microRNA miR-223 has a super-enhancer over the gene that extends far upstream. (A) *hsa-miR-223* and its gene neighborhood (chrX:65,203,177-65,260,919). (B) Tissue expression (bar graphs) from GTEx with selected mean TPM indicated. The thick horizontal line shows the region used for TPM determination. (C) RNA-seq for cell cultures is given for the first track as overlaid color-coded signal from myoblasts and lung fibroblasts and, for the second track, the data are just for the K562 cells. (D), (E), (F) and (G) and highlighting are as in Fig. 2. Blue dashed line, super-enhancer in monocytes and neutrophils; PFC, prefrontal cortex of brain. A second sample of monocytes from males (not shown) gave a similar chromatin state segmentation profile as for the exhibited female sample.

Less extensive, traditional enhancer chromatin was observed in T cells, B cells, and a B-cell derived LCL (GM12878; Fig. 3D and F, blue highlighting). The LCL as well as the hematopoietic cancer-derived K562 cell line preferentially express the miR-223 miRNA precursor with especially high levels in K562 cells, which also display a super-enhancer spanning the gene (Fig. 3C). A novel long noncoding RNA (lncRNA; Fig. 3C, orange line) as well as *hsa-mir-223* is strongly expressed from the K562 super-enhancer. This lncRNA is likely to be associated with the K562 enhancer activity but is stable, unidirectional (plus-strand RNA-seq, data not shown) and produced in considerable amounts unlike the typical enhancer-associated RNA (eRNA), which is short, bidirectional, and difficult to detect due to instability [44]. The LCL displayed none of this lncRNA, had much less extensive enhancer chromatin, and had much less TF binding in the distal region than did K562 cells (Fig. 3F).

### 3.3 Fibrosis-associated MMP9 and MMP2 genes display very different tissue-specific expression and epigenetic profiles

*MMP2* and *MMP9*, paralogous genes that encode related matrix metalloproteinases and are critically involved in cardiac fibrosis and cardiovascular disease [5], have very different tissue specificities. The concentrations of their respective processed protein products must be carefully balanced with levels of tissue inhibitors of metalloproteinases (TIMPs). *MMP9* is expressed preferentially in leukocytes (Fig. 1B). In contrast, *MMP2* is highly expressed in a broad spectrum of tissues, *e.g*., skin, aorta, lung, and adipose, and in many progenitor cell types, including skin and lung fibroblasts (Fig. 1C, 4; Supplemental Table S1). Curiously, *MMP2* as well as *AGT* and *CCN2/CTGF* displayed much higher expression in the right atrial appendage than in the left ventricle (Fig. 1), the only two regions of heart with expression data in the GTEx database. From the multiple *MMP2* isoforms containing different TSSs, 5’ Cap Analysis of Gene Expression (CAGE [31]) indicated that one predominant TSS is used in the examined cell cultures (Fig. 4B, broken arrow). Curiously, the 3’ end of *MMP9* overlaps the 3’ end of *SLC12A5-AS1*, a gene encoding a lncRNA whose 5’ end overlaps a brain-specific gene *SLC12A5*. However, this lncRNA gene, which has overall low expression, has an expression profile that is a mixture of that of *MMP9* and *SLC12A5* but more like that of *MMP9* [30]. Therefore, this lncRNA might help regulate both protein-coding genes that it overlaps.

**Fig. 4.**
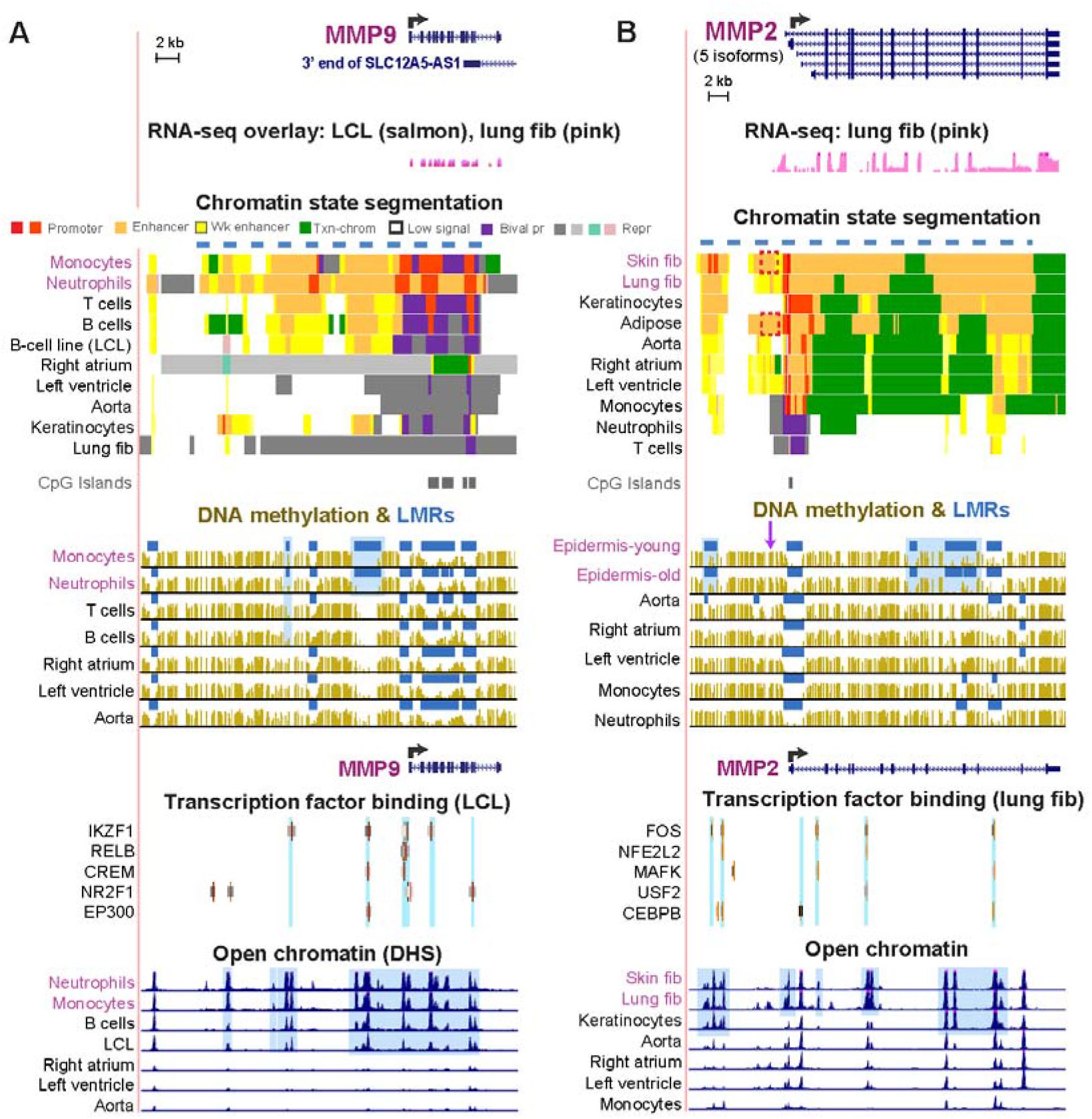
*MMP9* has a super-enhancer in neutrophils while *MMP2* has one in skin and lung fibroblasts. (A) *MMP9* (chr20:44,615,045-44,646,541) and (B) *MMP2* (chr16:55,503,063-55,541,196) as in Figs. 2 and 3. Blue highlighting, LMRs or DHS that are specific for monocytes/neutrophils or leukocytes; light blue highlighting for TF binding, TFs binding to DHS that are specific for leukocytes or fibroblasts/keratinocytes. Fib, fibroblast; bival pr, bivalent (repressed or weak) promoter chromatin containing H3K4me3 and H3K27me3.

We found super-enhancers in neutrophils and skin fibroblasts/lung fibroblasts, for *MMP9* and *MMP2*, respectively (Fig. 4A and B, dashed lines over the chromatin state tracks), which are likely to drive the high expression of these genes in these cell types. *MMP9* displayed more DNA hypomethylation and stronger DHS in monocytes and neutrophils than in B cells or the B-cell LCL (Fig. 4A, blue highlighting) consistent with cell type-specific differences in the extent of enhancer chromatin. Available TF-binding profiles for *MMP9* in the LCL and *MMP2* in a lung fibroblast cell strain showed that most of the TF binding was again in expression-associated DHS (Fig. 4, light blue highlighting). In the case of *MMP2*, tissue-specific DNA hypomethylation was most prominent in skin epidermis (for which no chromatin state profile is available). This could be related to the high level of *MMP2* expression in skin tissue (epidermal plus dermal layers; Fig. 1C). Although skin and aorta both exhibit very high levels of *MMP2* expression, skin displayed much more DNA hypomethylation than aorta, suggesting that different epigenetic strategies are sometimes used for maintaining similar levels of tissue-specific expression of a given gene (Figs. 1C and 4B). Heart failure-linked DNA hypomethylation was previously found [25] near *MMP2* and *CCN2* (Figures 4B and 5B, purple arrows), as discussed below.

**Fig. 5.**
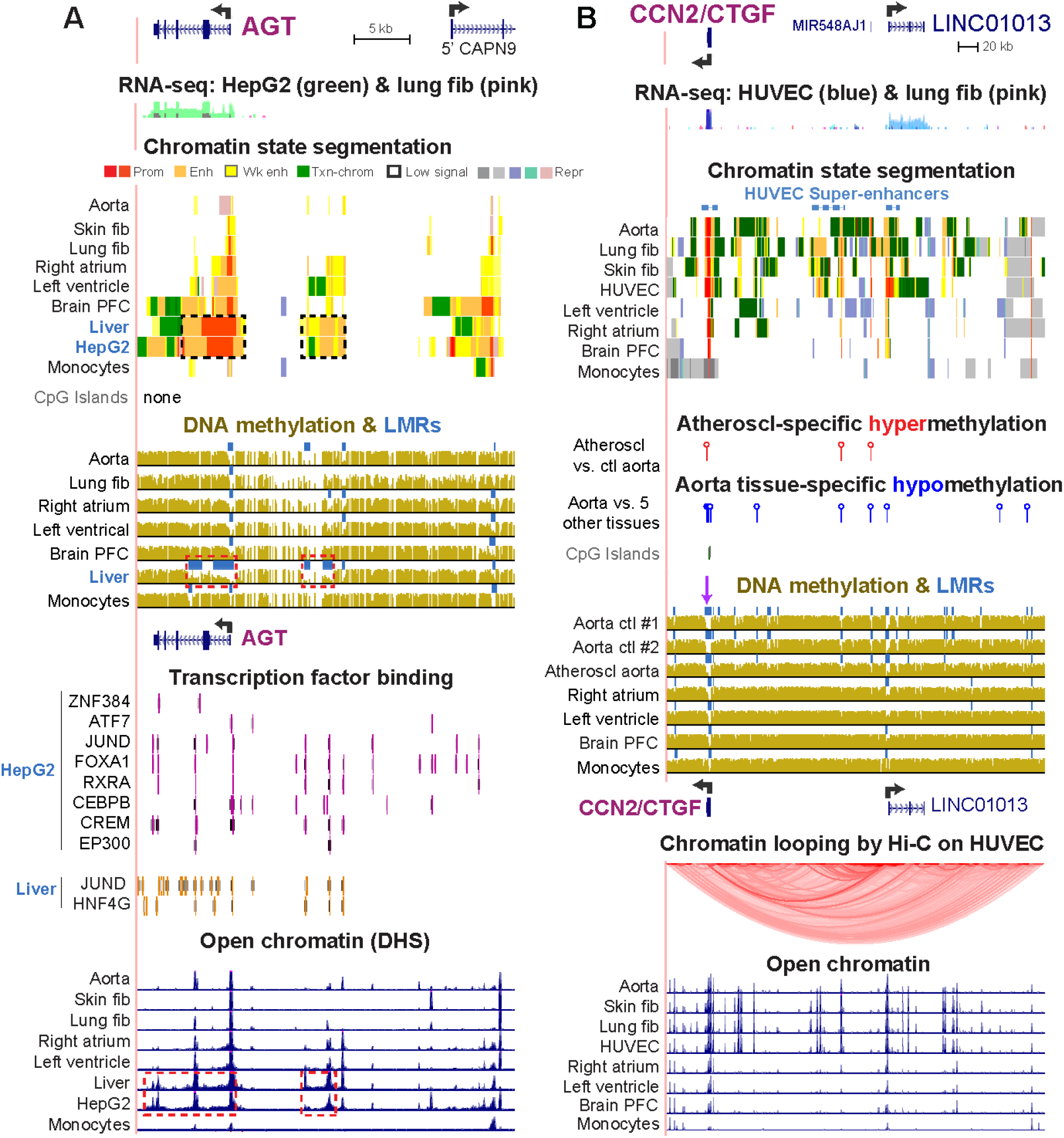
Enhancers in neighboring genes that are associated with expression in *AGT* or *CCN2/CTGF*. (A) *AGT* (chr1:230,835,715-230,892,908) and (B) *CCN2/CTGF* (chr6:132228520-132613558) as in Figs. 2 – 4 with the following additions. Significant differentially methylated regions (DMRs) in aorta vs. five other tissues (heart, skeletal muscle, lung, adipose or monocytes) and in an atherosclerotic (Atheroscl) aorta sample vs. three control (Ctl) aorta samples were determined for *CCN2* from bisulfite-seq data; blue, hypomethylated DMR; red, hypermethylated DMR. In addition, a chromatin looping profile [31, 45] (Hi-C technique) is shown for umbilical vein endothelial cells (HUVEC).

### 3.4 The liver-specific expression of AGT matches tissue-specific profiles of promoter and enhancer chromatin, open chromatin, and DNA methylation

*AGT*, a gene with critical functions in regulating blood pressure and vascular remodeling, is expressed highest in liver (Fig. 1E). Accordingly, liver and HepG2 cells displayed the most intragenic and intergenic promoter and enhancer chromatin and DNA hypomethylation as well as much TF binding (Fig. 5A, dotted boxes). Brain and heart, which had high levels of *AGT* RNA, although not as high as liver (Fig. 1E), exhibited considerable intragenic and intergenic *AGT* enhancer chromatin but less than for liver (Fig. 5A). In brain, liver, and HepG2, there was additional enhancer and promoter chromatin overlapping the gene body of *AGT*’s nearest upstream neighbor *CAPN9*, which encodes a cysteine protease. The enhancer profile at *CAPN9* paralleled the strong expression of *AGT* in these tissues (Fig. 1) but was at odds with the negligible expression of *CAPN9* [30] in these samples and did not correlate with expression of any other nearby gene. Therefore, we propose that this enhancer chromatin at *CAPN9* in liver and brain is upregulating its neighbor, *AGT*, rather than itself.

Previous studies revealed enhancer activity or enhancer chromatin only in part of the body of the *AGT* gene. Both an 80-bp associated enhancer element in the last exon of *AGT* that was active in HepG2 cells and a 1.4-kb androgen-responsive enhancer in intron 2 of *AGT* were described [28, 29]. The involvement of even more enhancer chromatin in regulating expression of *AGT* in liver and HepG2 cells is indicated not only by our description of their epigenomic profiles above, but also, by the many positively-acting TF binding sites throughout the gene and far upstream (Fig. 5A).

### 3.5 Preferential expression of CCN2/CTGF in aorta is accompanied by far-upstream DNA hypomethylation and enhancer chromatin over a cancer-associated ncRNA gene (LINC01013)

*CCN2/CTGF*, which is involved in many aspects of development (including angiogenesis) and homeostasis as well as in diseases (*e.g.*, myocardial infarction, atherosclerosis, hypertension, and cancer [22, 23]) had a very high median level of RNA (TPM, 2043) in aorta. Most examined cell cultures, including skin fibroblasts, lung fibroblasts, chondrocytes, umbilical vein endothelial cells (HUVEC) and HepG2 cells highly express this gene and displayed multiple regions of enhancer chromatin within a 350-kb region upstream of *CCN2* that includes several ncRNA genes (Figs. 1D and 5B; Supplemental Figs. S3 and S4). Many TFs colocalized with cell type-specific DHS in this region. HUVEC, which had the highest expression of *CCN2* among examined cell cultures (Supplemental Table S1), displayed the highest expression of *LINC01013*, a little-studied ncRNA gene in its far upstream intergenic region (Fig. 5B and Supplemental Fig. S4E). Coordinate expression of *LINC01013* and *CCN2* was previously observed in HUVEC after stimulation of the angiogenesis program genes *in vitro* with VEGFA [46]. The only other cell type expressing this ncRNA was chondrocytes. Little else is known about *LINC01013* except that it can repress chondrogenesis [47], and is associated with cancer progression [48]. Profiles of higher-order looping domains (topologically associating domains, TADs) in this neighborhood gave evidence for a loop between *CCN2* and the immediate upstream region of *LINC01013* in HUVEC (Fig. 5B, Hi-C [45]), but not in an LCL or a keratinocyte cell line (data not shown), which have negligible expression of *LINC01013* (Supplemental Table S1). Therefore, the evidence supports the hypothesis that transcription of *LINC01013* upregulates *CCN2* in HUVEC. However, coordinate expression between *CCN2* and *LINC01013* was seen only in HUVEC and chondrocytes and not in any tissue, including aorta. Therefore, any linkage of expression of these two genes is restricted to only some *CCN2*-expressing cell types.

The very high expression of *CCN2* in aorta was accompanied by less enhancer chromatin and weaker DHS in this region than seen in HUVEC or lung fibroblasts. HUVEC displayed three super-enhancers in this gene neighborhood, over *CCN2*, *LINC01013*, and an intergenic region in between them (Fig. 5B, dotted blue lines), but no super-enhancer was found in aorta. In interpreting the relationship between expression of this gene and the enhancer chromatin based upon GTEx RNA-seq data, we considered the possibility that *CCN2*/*CTGF* expression in GTEx samples was biased upward because ~38% of the hundreds of GTEx aorta samples were from individuals of > 60 yo. In contrast, the donor for the aorta chromatin state and DNA methylation analysis and for many other Roadmap samples [6] was a 34-yo male. However, this was not a problematic comparison because when we stratified GTEx data for expression of *CCN2*/*CTGF* aorta by age, the median values for expression of *CCN2/CTGF* for 59 donors between 20 and 49 yo were not significantly different from that for 42 donors between 50 and 79 yo.

One of the biggest differences between the epigenetics of the *CCN2* gene neighborhood in aorta and in other less highly expressing tissues was the presence of aorta-associated DNA hypomethylation. There were many aorta-specific LMRs at *CCN2* or far upstream of the gene that showed statistically significant DNA hypomethylation in aorta vs. the set of heart, lung, skeletal muscle, and monocyte samples (Fig. 5B, blue lollipops). Several of these LMRs overlapped aorta/HUVEC/fibroblast-associated DHS. Bisulfite-seq DNA methylation profiles were available for two samples from an 88-yo female [33] and one from the 34-yo Roadmap control aorta (Fig. 5B, Aorta ctl #1 [6]). One of the 88 yo donor’s samples was normal-appearing thoracic aorta (Aorta ctl #2) and the other from the same donor was a section of atherosclerotic aortic arch (Atheroscl aorta). Similar distinctive patterns of DNA hypomethylation were seen in all examined aorta samples (Fig. 5B, blue lollipops) but there was a small but significant amount of DNA hypermethylation in the atherosclerotic aorta vs. the control aorta samples in the vicinity of *CCN2* or *LINC01013* (Fig. 5B, red lollipops). Similarly, we and Zaina *et al.* [33, 34] previously showed that throughout the genome there were many more hypermethylated than hypomethylated DNA regions in atherosclerotic vs. control aortas. It remains to be determined if there are decreases in enhancer or promoter chromatin in some atherosclerosis-associated hypermethylated DNA regions.

## 4. Discussion

The fibrosis-related protein-coding genes that we studied, *NLRP3, MMP2, MMP9, CCN2/CTGF*, and *AGT*, are upregulated in diverse diseases. These include cardiac fibrosis and atherosclerosis [18, 19, 23], liver and lung fibrosis [23, 26, 49], rheumatoid arthritis and osteoarthritis [26], many types of cancer [23, 24, 50], and age-related increased susceptibility to vascular disease [51] and to lethality from COVID-19 [2]. The precise control of transcription of these genes is important because they can participate in restorative as well as pathological repair [52] or inflammation [18]. Age-related disease susceptibility to fibrosis and fibrosis-associated cardiovascular disease can be driven, in part, by age-related build-up of epigenetic mistakes in key genes [8]. Some of these mistakes might help abnormally upregulate expression of the studied genes, which are usually overexpressed rather than downregulated in disease. Despite the central role of epigenetics and, in particular, enhancers to gene regulation [38, 44], previous studies of the epigenetics of the genes that were highlighted in this study have been limited.

By using diverse whole-genome epigenetic profiles rather than just data from locus-specific studies [6], we were able to extend previous descriptions [25, 28, 29, 53] of transcription-regulatory chromatin at *AGT, CCN2/CTGF*, and *MMP2* and describe for the first time enhancer chromatin for *NLRP3.* NLRP3 protein is crucial for innate immunity and inflammation and is a major player in the induction of fibrosis [18]. Our results indicate that the examined cardiac fibrosis-associated genes, *NLRP3, MMP2, MMP9, CCN2/CTGF*, and the *NLRP3* mRNA-regulatory *hsa-miR-223*, have previously undescribed super-enhancers (clusters of traditional enhancers) containing foci of DNA hypomethylation, open chromatin, and TF binding that are very likely to help drive the high tissue-specific expression of these genes. Super-enhancers at other genes have been implicated in cardiovascular disease [11, 54]. *AGT*, the other fibrosis-related gene that we studied, has traditional enhancer chromatin as well as elongated promoter chromatin that can account for its very high expression in liver. Although enhancers are often thought of as upstream or downstream of genes, all six genes contained some of their enhancer chromatin in the gene body. Such gene-body enhancer regions might facilitate rapid transcription elongation as well as potentiate promoters, their better known role [55].

A caveat in our study is that our comparison of epigenetic and transcription profiles relies on steady-state mRNA levels, which depend, in part, upon factors affecting the stability of the RNA and, thereby, the measured mRNA levels. However, miRNA-targeted degradation usually only fine-tunes target mRNA levels, typically about two-fold [56]. Nonetheless, such fine-tuning can have important biological consequences, and more than one miRNA can target an mRNA. miR-223, which is encoded by the *hsa-miR-223* gene (Fig. 3), targets *NLRP3* mRNA as well as mRNAs for inflammation-related signaling proteins and TFs, downregulates healing of slow-healing wounds, upregulates granulopoiesis, downmodulates monocytic/macrophage differentiation, regulates the innate immune response in chronic obstructive pulmonary disease and asthma, and can affect cell viability, proliferation, and invasion in carcinogenesis [57–59].

The microRNA miR-223 as well as TTP, an RNA-binding protein that reduces levels of *NLRP3* mRNA, specifically target a long-form polyadenylation variant and not a short-form variant with a shorter 3’ untranslated region [60]. Importantly, there are similar amounts of both *NLRP3* RNA isoforms in macrophages, which we did not study because of a lack of available epigenome profiles, but the short-form variant is the predominant one in monocytes [60]. As monocytes differentiate to macrophages, there are decreases in miR-223 and increases in NLRP3 protein due to increased translation despite decreases in steady-state levels of *NLRP3* mRNA [43]. We found that *NLRP3* mRNA and the pri-miRNA precursor of miR-223 were most highly expressed in leukocytes, and the genes for both had super-enhancers specifically in monocytes and neutrophils (Figs. 1 - 3). The upregulation of *NLRP3* transcription as well as the downregulation of the activity of many *NLRP3*-targeting miRNAs, including miR-223, drive inflammasome activation in neutrophils and monocytes/macrophages in response to a priming trigger [27, 36, 43, 59]. Surprisingly, miR-223 is much more highly expressed in leukocytes than is *NLRP3*. These findings underscore the importance to NLRP3 inflammasome formation of posttranscriptional downregulation of miR-223 activity, probably by sponge or competing endogenous RNAs such as one that has been described to inhibit another mRNA target of miR-223 [61]. An additional important regulator of NLRP3 inflammasome formation involves direct modification of the NLRP3 polypeptide by ubiquitylation, sumoylation, and phosphorylation to control NLRP3 protein levels and activity [62]. The multiplicity of controls on this key inflammasome-forming protein indicates how important its precise levels are for its roles in innate immunity, inflammation, and disease [18, 62].

Our findings further elucidate previously reported disease-related epigenetic changes in several of the studied genes. A subregion 1.4 kb upstream of the distal TSS of *MMP2* was shown by Glezeva *et al.* [25] to be hypomethylated in left ventricle samples from heart-failure patients relative to analogous control samples. A disease-linked increase (almost 3-fold) in expression of *MMP2* accompanied this hypomethylation. Glezeva *et al.* had notated this 0.5-kb region as one without regulatory function. We found that this region (Fig. 4B, purple arrow) overlapped strong enhancer chromatin in skin fibroblasts and adipose tissue, both of which highly express this gene (Fig. 4B, dotted boxes). Therefore, our analysis of tissue-specific epigenetics in this region supports the importance of their heart-disease related findings about *MMP2.* Glezeva *et al.* [25] also found that heart failure was associated with significant DNA hypomethylation in a 1-kb subregion 0.3 kb downstream of the 3’ end of *CCN2/CTGF* and ~ 3-fold upregulation of this gene’s expression. We found that this DNA region is highly methylated and does not overlap enhancer or promoter chromatin in heart samples. In contrast, it is mostly unmethylated and overlaps promoter or enhancer chromatin and expression-linked peaks of open chromatin in lung and skin fibroblasts and aorta (Supplemental Fig. S3, purple arrow). Therefore, the disease-associated hypomethylation at *CCN2* in heart failure described by Glezava et al. [25] might signify disease-related enhancer formation downstream of *CCN2*.

Despite only limited amounts of genome-wide data being available for TF binding, we found that some important TFs or cofactors bound to transcription regulatory elements in cells expressing these genes. For example, *CTGF*, whose promoter binds the histone acetyltransferase coactivator EP300 (p300) in response to TGFβ signaling in mouse fibroblasts [63], also binds this protein at far upstream and downstream enhancer chromatin within open chromatin in HepG2 cells (Supplemental Fig. S3). Therefore, the effects of TGFβ signaling on *CTGF* expression probably involve enhancer as well as promoter upregulation mediated by EP300. Similarly, many studies show that NFκB is important for induction of transcription of *NLRP3* in response to pathogen-associated molecular patterns but reports of NFκB binding to the *NLRP3* gene region have described only the *NLRP3* promoter and immediate upstream region [36, 49]. We found multiple NFκB binding sites in regions of open chromatin in the body of the *NLRP3* gene, far upstream, as well as in the promoter region in a B-cell LCL (Fig. 2D and E), which expresses this gene only weakly. Importantly, these sites displayed much stronger peaks of open chromatin in *NLRP3*-highly expressing monocytes and neutrophils. Therefore, it is very likely that there are yet more and stronger TF binding sites in these myeloid cells. The examined liver samples in this study had barely detectable expression of *NLRP3* RNA, no *NLRP3*-associated enhancer chromatin, and only some promoter chromatin at the TSS (Fig. 2). However, *NLRP3* expression can be induced in liver upon inflammatory stimulation [49]. We predict that this induction will involve the acquisition of monocyte/macrophage-like intragenic and intergenic enhancer chromatin. A comparison of *NLRP3* enhancer/promoter chromatin vs. expression levels in various tissues (Figs. 1 and 2) showed that amounts of enhancer chromatin were a better predictor of expression than were amounts of promoter chromatin.

Our study shows the importance of extending epigenetic studies of pathogenic samples to include intragenic and far distal enhancers as well as super-enhancers. By examining the epigenetics of whole gene neighborhoods, we obtained novel insights into regulation of these genes during normal development. For example, our study implicates enhancer chromatin overlapping *CAPN9* in liver and brain in the upregulation of *AGT, CAPN9*’s downstream neighbor. *CAPN9* itself has undetectable expression in these tissues. In addition, *CCN2/CTGF* and its cancer-related neighbor ncRNA gene *LINC01013* [48], which is 182 kb upstream, provide another likely example of long-distance transcription-regulatory interactions spanning genes but only in HUVEC and chondrocytes. It was previously shown that this gene/lincRNA gene pair exhibit parallel changes in expression after exposure of HUVEC to VEGFA [46]. These types of long-distance interactions in *cis*, our finding of super-enhancers at most of the studied genes, and the dense clusters of TF binding sites that we describe at these super-enhancers provide new insights that can empower the development of drugs aimed at modulating abnormal expression of these important fibrosis-related genes.

## Supporting information

Chandra et al_Supplemental Figures and Table_CardiacFibrosis

## Acknowledgements

This research was supported in part using high performance computing (HPC) services provided by Information Technology at Tulane University, New Orleans, LA.

