## Supplementary material for "Epigenetics and expression of key genes associated with cardiac fibrosis: *NLRP3, MMP2, MMP9, CCN2/CTGF*, and *AGT*": Chandra et al_Supplemental Figures and Table_CardiacFibrosis

Supplemental Figures S1 – S4

**Fig. S1.** ***NLRP3* is expressed at a high level in monocytes and a low level in B cells and a B-cell lymphoblastoid cell line**

**Fig. S2**. **Epigenetics of other genes that encode components of NLRP3 inflammasomes: *CASP1* and *PYCARD*, but not *NEK7*, display leukocyte-associated preferential expression and enhancers**

**Fig. S3.** **Transcription factor (TF) binding to the *CCN2/CTGF* gene neighborhood**

**Fig. S4. Non-coding RNA genes in the neighborhood of *CCN2/CTGF***

Table S1

**Supplemental Table S1. Expression of studied cardiac fibrosis-related genes in cell cultures**

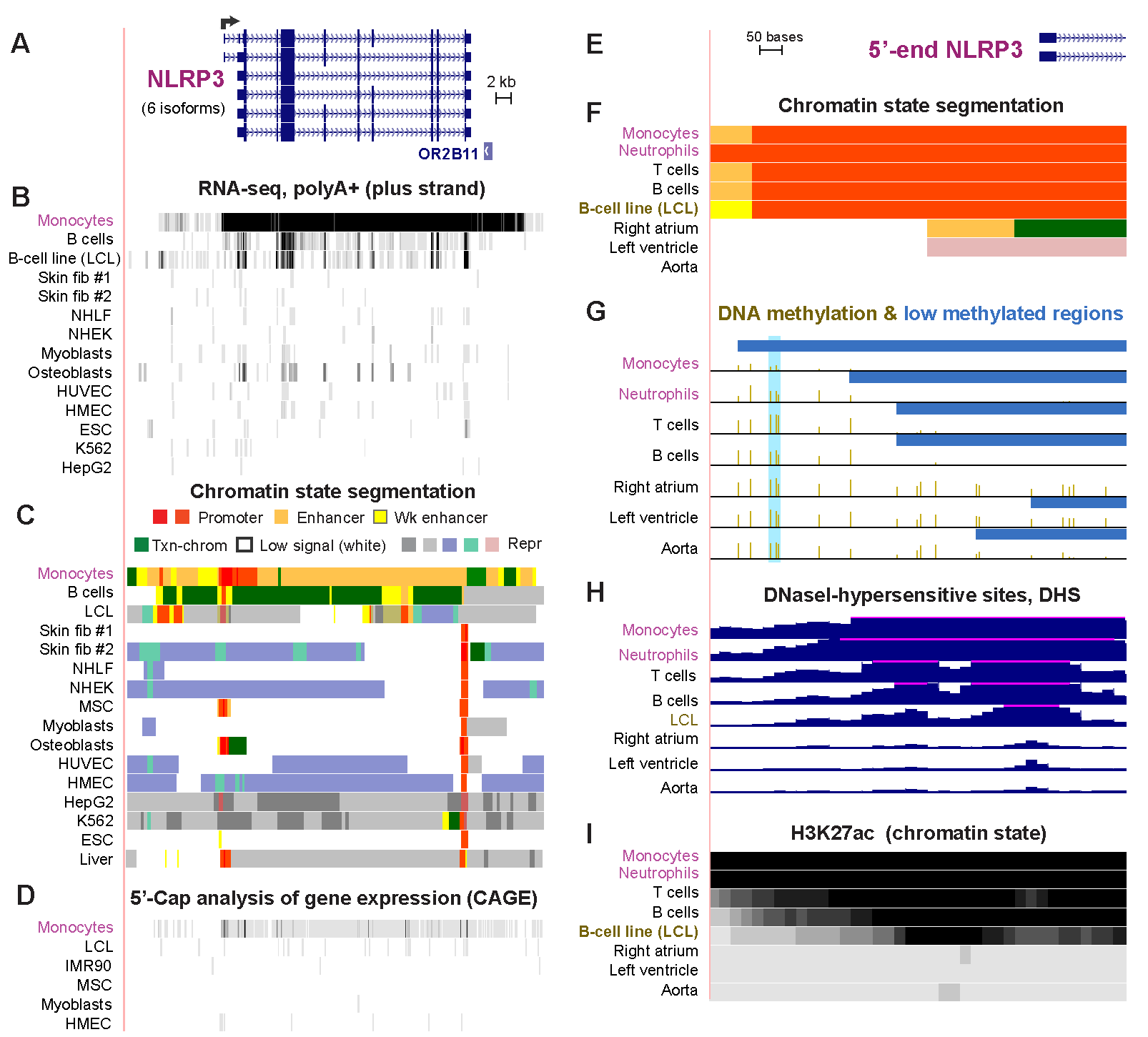

**Fig. S1.** ***NLRP3* is expressed at a high level in monocytes and a low level in B cells and a B-cell lymphoblastoid cell line.** (A) Six RefSeq isoforms of *NLRP3* (chr1:247,566,540-247,622,212), with the broken arrow indicating the TSS supported by data in Panels (B) – (D) as well as by the GTEx database <https://www.gtexportal.org>, which indicates predominant expression of the uppermost isoform. (B) RNA-seq data for the plus strand (polyA^+^ strand specific long (>200 nt) RNA; vertical viewing range of 0 – 15). (C) Chromatin state segmentation using the 18-state model (Roadmap Epigenomics Project) instead of the 25-state model as in Fig. 2; the two models use the same histone modification data but somewhat different algorithms for calculating the chromatin state. (D) 5’ Cap analysis of gene expression (CAGE, plus-strand signal; ENCODE Project, UCSC Genome Browser); only the indicated TSS aligns with a CAGE signal. (E) – (I) Zoomed-in view of the 5’ end of *NLRP3* and its promoter region (chr1:247,578,703-247,579,654) showing DNA methylation, bisulfite-seq; open chromatin, DNase-seq for open chromatin DNaseI-hypersensitive sites (DHS), and H3K27ac signal. Fib, fibroblasts; NHLF, lung fibroblasts; NHEK, epidermal keratinocyte; HUVEC, umbilical vein endothelial cells; HMEC, mammary epithelial cells; ESC, embryonic stem cells; skin fib #1 and #2, two independent skin fibroblast primary cell cultures; HepG2, hepatocellular carcinoma cell line; K562, myelogenous leukemia cell line; blue highlighting, a previously described DNA hypomethylated site (Wei et al., 2016, see the main text for the reference). All tracks are from the UCSC Genome Browser or its hubs and hg19.

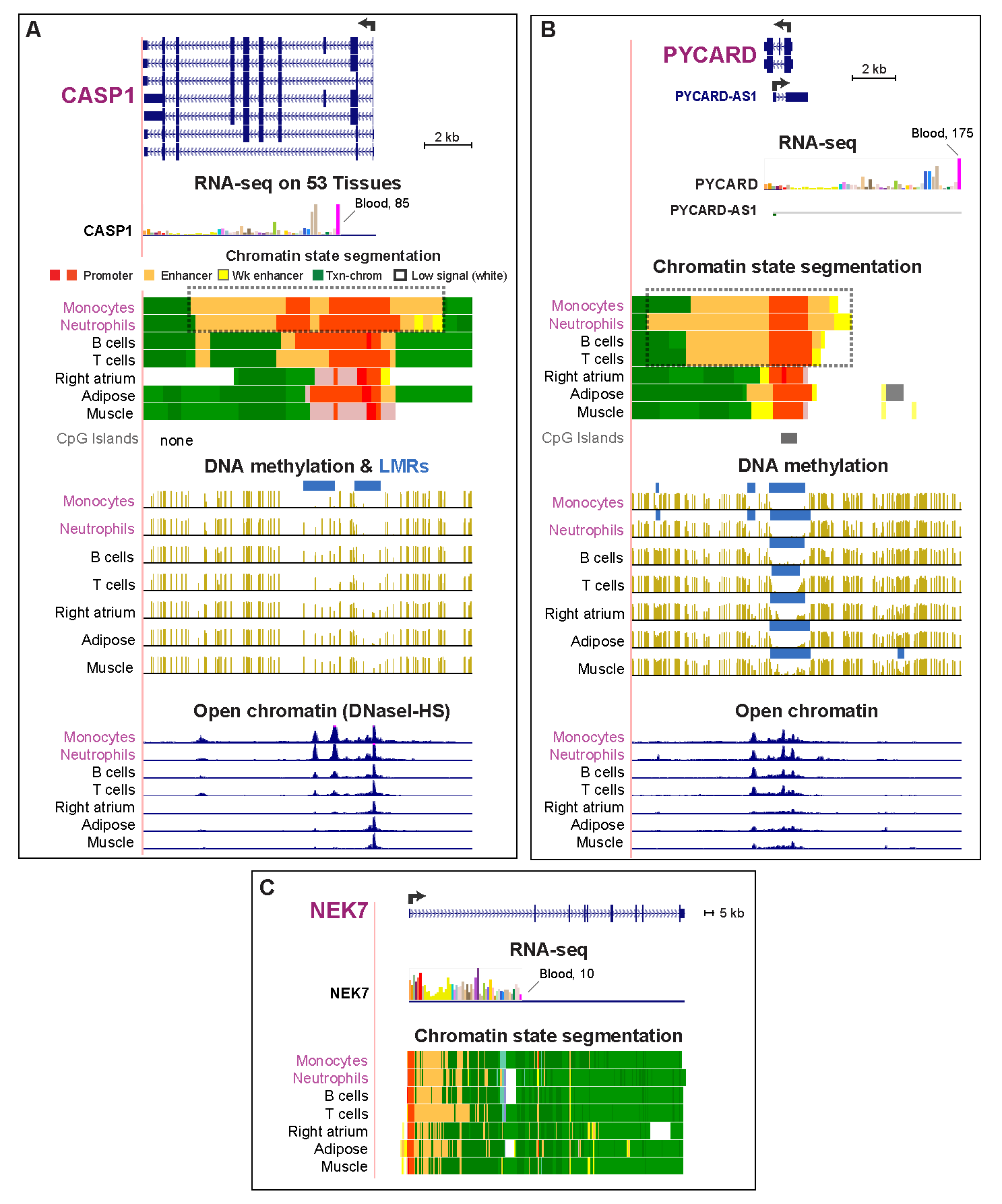

**Fig. S2**. **Epigenetics of other genes that encode components of NLRP3 inflammasomes: *CASP1* and *PYCARD*, but not *NEK7*, display leukocyte-associated preferential expression and enhancers**. (A) *CASP1* (chr11:104,896,136-104,910,02). (B) *PYCARD* (chr16:31,206,676-31,221,844). (C) *NEK7* (chr1:198,106,285-198,304,281). Epigenetic tracks are as described in Fig. S1 except that the 25-state model for chromatin state was used. Median RNA-seq levels (TPM) from hundreds of samples are shown as linear bar graphs in which the black vertical line indicates the area from which the mRNA signal was calculated.

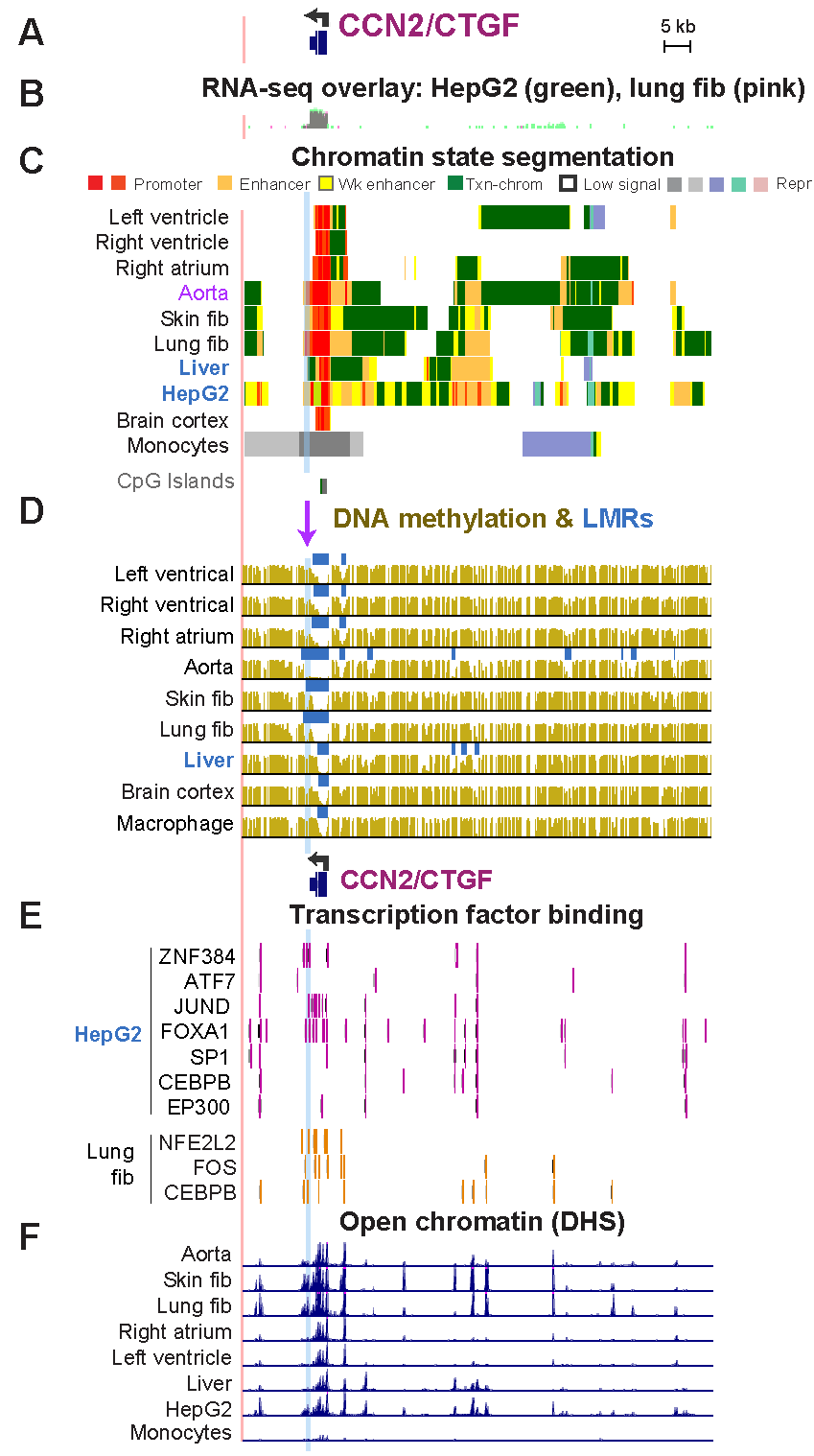

**Fig. S3.** **Transcription factor (TF) binding to the *CCN2/CTGF* gene neighborhood.** (A) *CCN2* and its upstream gene neighborhood (chr6:132,256,021-132,348,643). (B) Overlay RNA-seq tracks showing high expression of *CCN2* in HepG2 and lung fib. (C) Chromatin state segmentation tracks from the 18-state model. (D) Bisulfite-seq tracks for DNA methylation showing low methylated regions (LMRs). (E) TF binding sites in HepG2 cells or lung fibroblasts. (E) DNase-I hypersensitive sites (DHS). Blue highlighting and the purple arrow show the region of previously described DNA hypomethylation in left ventricle samples from patients with heart failure vs. controls (Glezeva et al., 2019, ref. 24, and see main text).

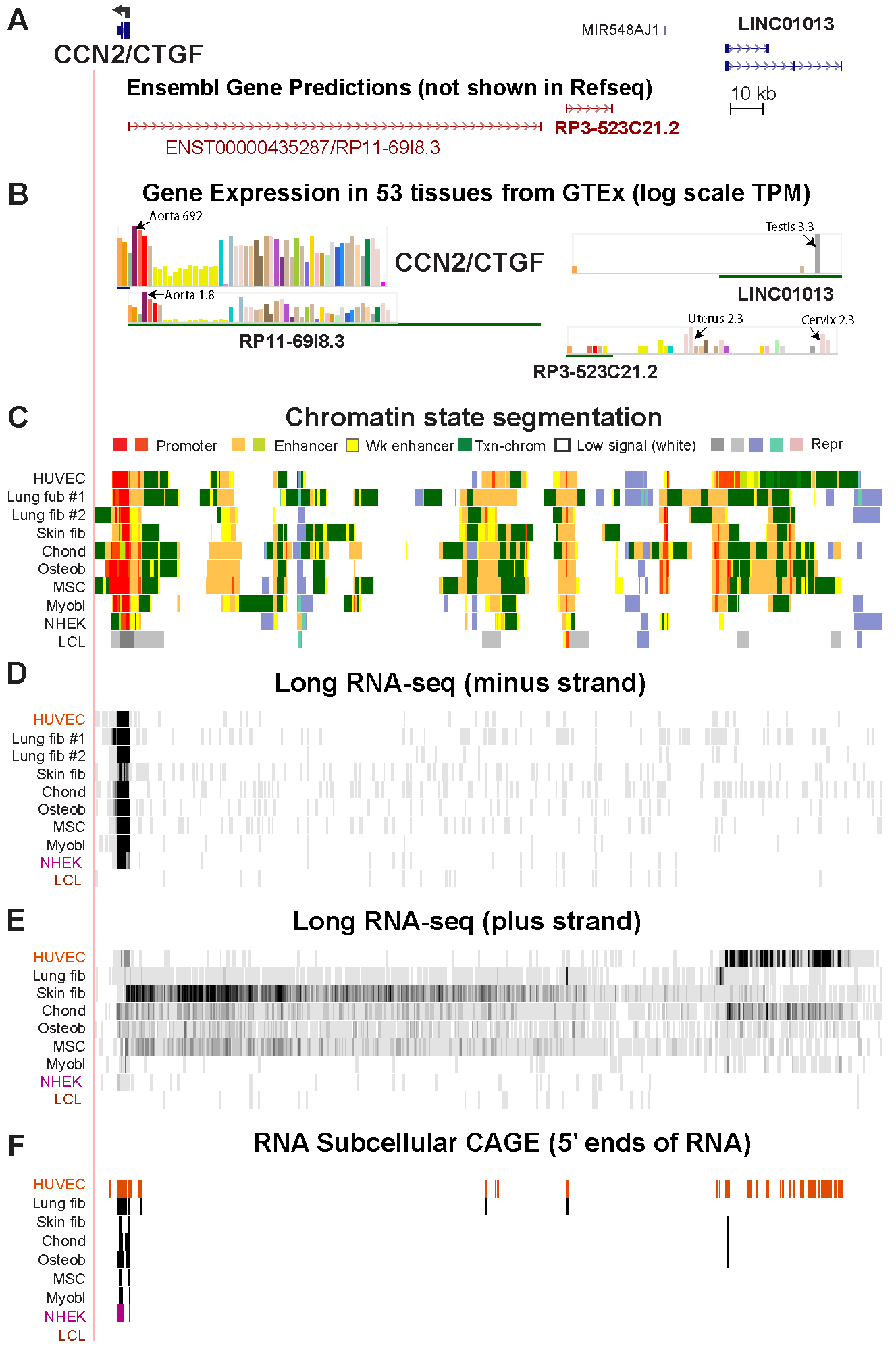

**Fig. S4. Non-coding RNA genes in the neighborhood of *CCN2/CTGF.*** (A) ncRNA genes in the upstream gene neighborhood of *CCN2* (chr6:132,262,203-132,502,620). (B) GTEx RNA-seq data showing high expression of *CCN2* and ncRNA gene *RP11-6918.3* in aorta. (C) Chromatin state segmentation tracks (18-state model) showing multiple regions of enhancer chromatin in this subregion in HUVEC, lung fib and skin fib. (D) RNA-seq data for the minus strand (vertical viewing range, 0 - 100) showing high expression of *CCN2* in most cell lines. (E) RNA-seq data for the plus strand (vertical viewing range, 0 - 30) showing high expression of *LINC01013* in HUVEC and chondrocytes. (F) 5’ Cap analysis of gene expression (CAGE, transcriptional start sites based on Hidden Markov Modeling for pooled replicates; ENCODE Project; UCSC Genome Browser). See Supplemental Table S1 for quantitation of RNA levels from non-strand specific RNA-seq, which is more accurate than strand-specific RNA-seq.

| **Supplemental Table S1. Expression of studied cardiac fibrosis-related genes in cell cultures^a^** | | | | |  |  |  |
| --- | --- | --- | --- | --- | --- | --- | --- |
|  |  | **RNA levels from RNA-seq (FPKM) from the ENCODE Project** | | | | | |
| Gene ID | Coordinates in hg19 | Myoblasts | Lymphoblastoid cell line (GM12878) | Embryonic stem cells (ESC) | HUVEC^b^ | NHEK^b^ | NHLF^b^ |
| NLRP3 | chr1:247579457-247612410 | 0.1 | 0.3 | 0.1 | 0.1 | 0.1 | 0.1 |
| MMP2 | chr16:55512750-55620578 | 231.1 | 0.0 | 22.6 | 304.8 | 29.5 | 276.9 |
| MMP9 | chr20:44637521-44688789 | 0.0 | 0.4 | 2.5 | 0.0 | 1.6 | 0.0 |
| AGT | chr1:230838268-230937749 | 3.7 | 0.0 | 0.0 | 0.1 | 0.0 | 1.1 |
| CCN2/CTGF | chr6:132129155-132398533 | 188.7 | 0.0 | 36.8 | 534.9 | 36.7 | 146.3 |
| LINC01013 (ENSG00000228495; ENST00000458028; RP3-523C21.1) | chr6:132453054-132490514 | 1.0 | 0.0 | 0.1 | 62.0 | 0.0 | 0.2 |
| ^a^RNA-seq data from non-strand specific analyses was obtained as described in Materials and Methods from cell cultures | | | | | | |  |
| ^b^HUVEC, human umbilical vein cells; NHEK, normal human epidermal keratinocytes; NHLF, normal human lung fibroblast, all of which were primary cultures | | | | | | | |
